# PCR-free library preparation greatly reduces stutter noise at short tandem repeats

**DOI:** 10.1101/043448

**Authors:** Melissa Gymrek

## Abstract

Over the past several decades, the forensic and population genetic communities have increasingly leveraged short tandem repeats (STRs) for a variety of applications. The advent of next-generation sequencing technologies and STR-specific bioninformatic tools has enabled the profiling of hundreds of thousands of STRs across the genome. Nonetheless, these genotypes remain error-prone, hindering their utility in downstream analyses. One of the primary drivers of STR genotyping errors are “stutter” artifacts arising during the PCR amplification step of library preparation that add or delete copies of the repeat unit in observed sequencing reads. Recently, Illumina developed the TruSeq PCR-free library preparation protocol which eliminates the PCR step and theoretically should reduce stutter error. Here, I compare two high coverage whole genome sequencing datasets prepared with and without the PCR-free protocol. I find that this protocol reduces the percent of reads due to stutter by more than four-fold and results in higher confidence STR genotypes. Notably, stutter at homopolymers was decreased by more than 6fold, making these previously inaccessible loci amenable to STR calling. This technological improvement shows good promise for significantly increasing the feasibility of obtaining high quality STR genotypes from next-generation sequencing technologies.

## 1 Introduction

Short tandem repeats (STRs) are repetitive tracts of DNA comprised of an underlying 1-6 pair motif. Their repetitive structure induces DNA polymerase slippage events that add or delete copies of the repeat unit, resulting in exceptionally high mutation rates. This same biological process presumably leads to errors in the number of repeats that accumulate during the PCR amplification step used in library preparation protocols for next-generation sequencing. As a result, sequencing data often contains errors in the number of repeats, referred to as “PCR stutter”, complicating STR genotyping from short reads.

Figure 1 gives an example of a sequencing error likely due to PCR stutter. This example is from a locus on the X chromosome from a male sample, where there is expected to be only one true allele. One read is off by a single repeat unit (missing an AGAT) from the presumed true allele. This problem becomes even more complicated at diploid loci that may have two true alleles with separate stutter noise patterns, making it difficult to determine the true genotype. Recently, we developed lobSTR [4], an algorithm for STR calling that attempts to model PCR stutter noise when scoring genotypes. However, PCR stutter remains one of the major sources of error at STRs.

**Figure 1:**
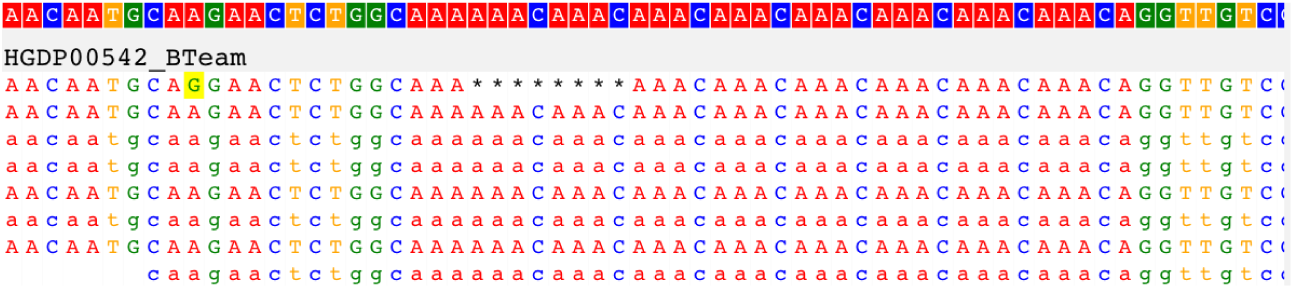
Example of PCR stutter error. Alignment visualization was produced by PyBamView [3].

Illumina recently developed the TruSeq “PCR-free” library preparation protocol [2], which eliminates the PCR amplification step from traditional library preparation techniques. I and others have hypothesized that this protocol may dramatically reduce the PCR stutter issue at STRs. Below I compare two whole genome sequencing datasets of similar quality prepared with and without PCR, and show evidence that the PCR free protocol reduces the stutter rate by more than four fold.

## 2 Methods

### 2.1 Datasets

We compared stutter noise in whole genome sequencing datasets for two samples. Sample HGDP00542 from Papua New Guinea was sequenced to 25.9x coverage as part of a pooled sequencing run as reported in Meyer *et al.* [6]. Library preparation was performed as described by Rohland and Reich [7] and included a PCR enrichment step. The second sample is for a Pakistani individual (sample HGDP00208) sequenced to 37.6x coverage following Illumina’s TruSeq PCR-free library preparation protocol. This sample was sequenced as part of the Simons Genome Diversity Project described at [1].

Both datasets consist of 100bp paired end reads for male samples sequenced on Illumina HiSeq2000 sequencers. Besides the difference in library preparation protocols, these datasets are expected to have similar quality.

### 2.2 lobSTR commands

STR genotypes were obtained using lobSTR [4], a specialized algorithm for profiling STR variation from high-throughput sequencing data. BAM files were first sorted by read name using samtools [5]. They were then aligned to STR containing regions using lobSTR v2.0.4. Genotypes were generated using the lobSTR allelotype tool v2.0.5. The specific commands used to process each BAM file are given in the Supplemental Methods.

### 2.3 Measuring stutter noise

For each sample, we estimated stutter noise at loci on the X chromosome. Since both samples are male, X chromosome calls are expected to be hemizygous and therefore haploid. Reads not matching the true allele could result from alignment artifacts or stutter noise. Here we assume minimal impact from alignment errors. We obtained all X chromosome calls made using five or more informative reads and with a unique modal allele (the allele supported by the most reads) assumed to represent the true allele. Any read not matching the “true” allele was deemed a stutter error.

## 3 Results

### 3.1 The PCR-free dataset has 4-fold fewer stutter reads

We first measured the percentage of reads determined to result from stutter noise. Table 1 gives the STR stutter rate broken down by motif size. The stutter rate clearly decreases with motif length, with homopolymers having by far the highest stutter rate. On average, the PCR free dataset had more than four-fold fewer percentage reads resulting from stutter. Note that the larger difference for hexanucleotide repeats is likely due to noise stemming from the small amount of loci analyzed.

**Table 1:** Comparison of STR stutter rates using PCR vs. PCR-free library preparation protocols.

| Motif length | Stutter prob. - with PCR | Stutter prob. - PCR free | Fold decrease |
| --- | --- | --- | --- |
| 1 | 0.17 | 0.027 | 6.30 |
| 2 | 0.038 | 0.011 | 3.45 |
| 3 | 0.011 | 0.0029 | 3.79 |
| 4 | 0.0069 | 0.0016 | 4.31 |
| 5 | 0.0074 | 0.0015 | 4.93 |
| 6 | 0.014 | 0.0010 | 14.0 |
| Average | 0.051 | 0.011 | 4.63 |

### 3.2 Stutter error sizes are similar across protocols

While the rate of stutter reads was greatly reduced in the PCR-free dataset, the distributions of stutter error sizes was similar. Figure 2 shows the distributions of error sizes for each motif length 2-5. Stutter noise tends to come in multiples of the repeat unit length with a strong bias toward decreasing the number of repeats of the true allele, matching what we have observed in previous work [4]. This general trend holds for both datasets, with the PCR-free dataset having a slightly lower downward bias.

**Figure 2:**
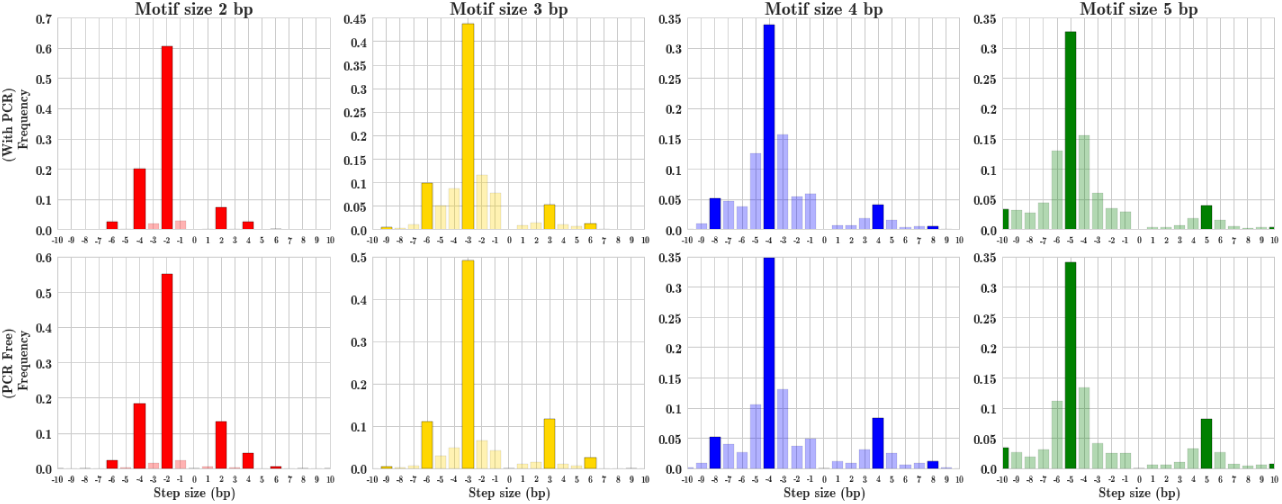
Stutter error size distributions by motif for each protocol. Top: with PCR. Bottom: without PCR.

### 3.3 PCR-free datasets show improved genotype quality scores

Finally, we determined whether the PCR-free protocol resulted in higher quality genotype calls by comparing the distributions of genotype quality scores. For each genotype call, lobSTR outputs a quality score defined as:

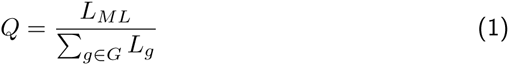

where *L*_*ML*_ is the likelihood of the maximum likelihood genotype, *G* is the set of all possible diploid genotypes, and *L*_*g*_ is the likelihood of genotype *g*. A *Q*-value close to 1 implies a confident call, whereas *Q* close to 0 means the call is not much better than the other possible genotype calls. We compared the average *Q* for different coverage levels across the two datasets, shown in Figure 3. Scores are significantly higher at all coverage levels in the PCR-free dataset.

**Figure 3:**
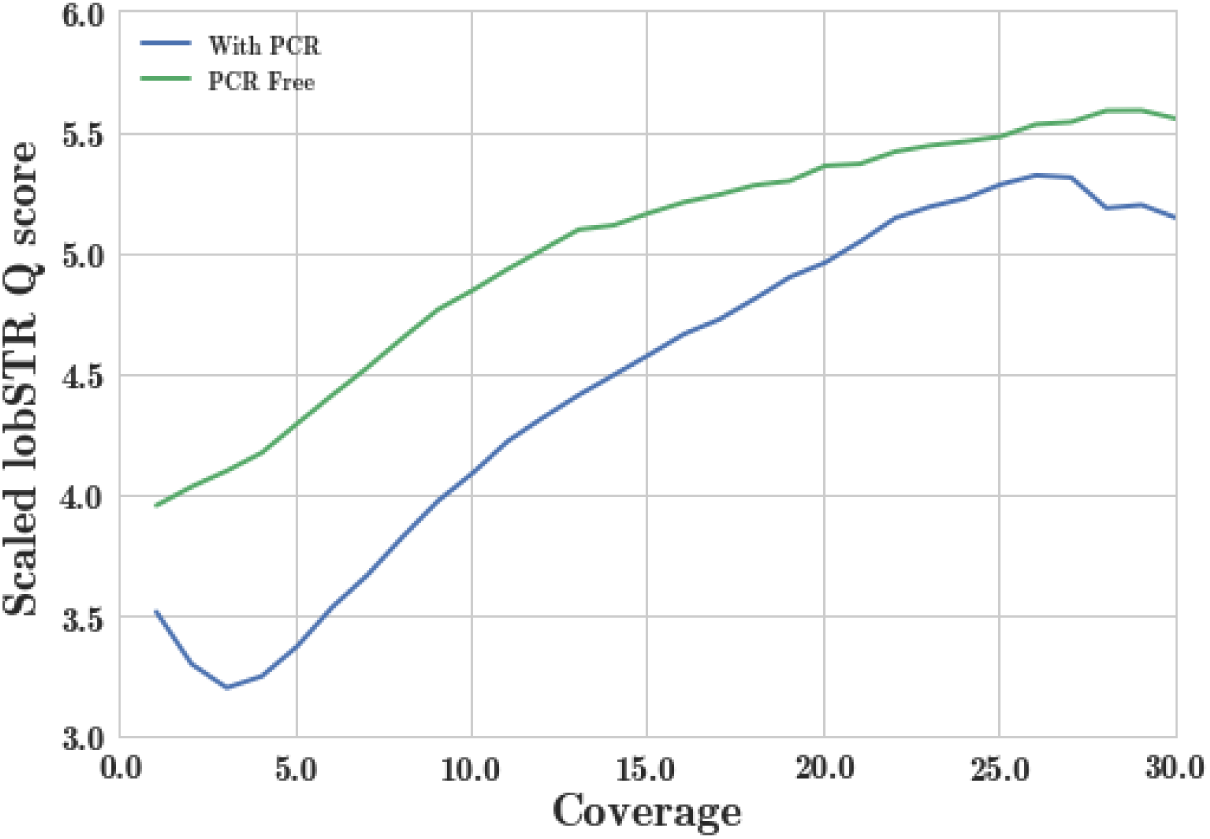
Average scaled *Q* score (—log_10_(1 — *Q*)) by coverage level for the PCR-free (blue) and with PCR (green) datasets.

## 4 Discussion

In conclusion, the Illumina PCR-free protocol has great promise for improving STR calls from high throughput sequencing datasets. While I only compared two datasets here, I see similar trends in other datasets: PCR-free samples have much lower stutter noise at STRs. Of particular note is the improvement at homopolymers, which have until now been nearly impossible to call with reasonable accuracy. STR call quality is likely to dramatically improve as more and more sequencing datasets are generated using this protocol.

## 5 URLs

- lobSTR software: http://lobstr.teamerlich.org/
- PyBamView software: mgymrek.github.io/pybamview
- Simons Genome Diversity Project: https://www.simonsfoundation.org/life-sciences/simons-genome-diversity-project-dataset/
- HGDP00542 BAM alignments http://cdna.eva.mpg.de/denisova/BAM/human/

## 6 Acknowledgements

The author would like to acknowledge Thomas Willems for his helpful comments on this manuscript.

## 7 Supplemental Methods

The following give the commands to process each BAM file using lobSTR.

~~~
# lobSTR alignment
~~~

~~~
lobSTR \
~~~

~~~
‐f ${BAM_SORTED_BY_NAME} \
~~~

~~~
‐‐bampair \
~~~

~~~
‐‐index-prefix resource_bundles/hg19/lobstr_v2.0.3_hg19_ref/lobSTR_ \
~~~

~~~
‐‐rg-sample ${SAMPLE} \
~~~

~~~
‐‐rg-lib ${SAMPLE} \
~~~

~~~
‐‐out ${SAMPLE} \
~~~

~~~
‐p 20 \
~~~

~~~
‐‐fft-window-size 16 \
~~~

~~~
‐‐fft-window-step 4 \
~~~

~~~
‐‐bwaq 15 \
~~~

~~~
‐v
~~~

~~~
samtools sort ${SAMPLE}.aligned.bam ${SAMPLE}.sorted
~~~

~~~
samtools index ${SAMPLE}.sorted.bam
~~~

~~~
# Train noise model
~~~

~~~
samtools view ‐b ${SAMPLE}.sorted.bam chrX > train_${SAMPLE}.sorted.bam
~~~

~~~
samtools index train_${SAMPLE}.sorted.bam
~~~

~~~
allelotype \
~~~

~~~
‐‐command train \
~~~

~~~
‐‐noise_model ${SAMPLE} \
~~~

~~~
‐‐index-prefix resource_bundles/hg19/lobstr_v2.0.3_hg19_ref/lobSTR_ \
~~~

~~~
‐‐strinfo resource_bundles/hg19/lobstr_v2.0.3_hg19_strinfo.tab \
~~~

~~~
‐‐haploid chrX \
~~~

~~~
‐‐bam train_${SAMPLE}.sorted.bam \
~~~

~~~
‐‐min-border 5 \
~~~

~~~
‐‐min-bp-before-indel 7 \
~~~

~~~
‐‐maximal-end-match 15 ‐v
~~~

~~~
# Run allelotype
~~~

~~~
allelotype \
~~~

~~~
‐‐command classify \
~~~

~~~
‐‐noise_model ${SAMPLE} \
~~~

~~~
‐‐index-prefix resource_bundles/hg19/lobstr_v2.0.3_hg19_ref/lobSTR_ \
~~~

~~~
‐‐strinfo resource_bundles/hg19/lobstr_v2.0.3_hg19_strinfo.tab \
~~~

~~~
‐‐bam train_${SAMPLE}.sorted.bam \
~~~

~~~
‐‐min-border 5 \
~~~

~~~
‐‐min-bp-before-indel 7 \
~~~

~~~
‐‐maximal-end-match 15 ‐v \
~~~

~~~
‐‐out ${SAMPLE}
~~~

~~~
vcf-sort ${SAMPLE}.vcf | bgzip ‐c > ${SAMPLE}.vcf.gz
~~~

~~~
tabix ‐p vcf ${SAMPLE}.vcf.gz
~~~

